# A correlative quantitative phase contrast and fluorescence super-resolution microscope for imaging molecules in their cellular context

**DOI:** 10.1101/2025.10.13.682186

**Authors:** Yujin Bao, Zach Marin, Xiongchao Chen, Qiong Liu, Yazgan Tuna, Sylvi Stoller, Florian Schueder, Chuyue Zhang, Suet Yin Sarah Fung, Min Wu, Karla M. Neugebauer, Jonathon Howard, Chi Liu, David Baddeley, Michael Shribak, Joerg Bewersdorf

## Abstract

Fluorescence microscopy has been widely used to reveal the spatial distribution of specifically labeled molecules, but it is blind to cellular context. Quantitative phase contrast microscopy (QPC) provides such complementary information. Here we have developed a platform that combines the QPC technique of correlative orientation-independent differential interference contrast (OI-DIC) microscopy with single-molecule super-resolution fluorescence microscopy. We demonstrate a detection sensitivity of 0.05 nm optical path difference, sufficient to detect single microtubules, and show its capability of 3D super-resolution fluorescence imaging in the cellular context. Additionally, we report deep-learning enabled digital staining, identifying nuclei, mitochondria and lipid droplets from OI-DIC data and demonstrate the potential of this approach for long-term live-cell imaging of organelles of interest without the need for fluorescence. OI-DIC can be easily integrated into most fluorescence microscopes and is readily adoptable by microscopy labs.

**ONE-SENTENCE TEASER:** A highly sensitive technique to visualize sub-cellular structures, dynamics, and molecules in their cellular context.

## Introduction

Understanding the inner workings of a cell and cellular interactions is a crucial component of biological studies. Among the methods available, fluorescence microscopy reveals the spatial distribution of specifically labeled molecules of interest. The development of super-resolution microscopy (SRM) in recent decades has overcome the diffraction limit (∼250 nm) and now enables the observation of nanoscale structures within cells. Among these methods, single-molecule localization microscopy (SMLM) (*1*) stands out for its <20-nm resolution and relatively simple implementation. SMLM instruments allow users to not only count molecules but also track moving molecules or other small particles and thereby observe dynamic phenomena such as the association and dissociation of molecules with specific targets.

However, fluorescence microscopy is blind to the cellular context the molecules are embedded in. Label-free microscopy methods, such as the well-established techniques of brightfield microscopy, Zernike phase contrast (PC) microscopy (*2*) and differential interference contrast (DIC) microscopy (*3*)(*4*), provide this complementary information. Since they require only very low light doses, they cause negligible phototoxicity and are therefore well-suited for long-term live-cell imaging. However, in contrast to fluorescence microscopy, the recorded signal of these classical label-free imaging methods at any pixel is not directly proportional to the number of molecules in the corresponding sample volume, making it difficult to extract quantitative information about the sample. To solve this problem, a range of quantitative phase contrast (QPC) microscopy techniques have been developed (*5*)(*6*). These methods measure the phase delay of light, which is proportional to the dry mass density (primarily proteins, lipids, and nucleic acids) at the corresponding location in the sample.

By combining the imaging modalities of fluorescence and QPC, specific molecular information can be directly correlated with general cellular information. For instance, diffraction phase microscopy (DPM) (*7*) was combined with epifluorescence to observe dry mass changes within subcellular regions indicated by fluorescent tags in live kidney cells (*8*). Furthermore, Spatial Light Interference Microscopy (SLIM) (*9*) was correlated with epifluorescence to study cell cycle-dependent growth (*10*) and with light-sheet microscopy to image oleaginous yeast in microfluidics (*11*). Another example is remote-focus Quantitative Label-free Imaging with Phase and Polarization (QLIPP) (*12*) correlated with oblique light-sheet microscopy to analyze cellular dynamics (*13*). QPC has also been combined with super-resolution microscopy techniques realizing ∼170 nm lateral resolution in the QPC channel (*14*, *15*) and ∼90 nm lateral resolution in the fluorescence channel (*14–17*). Additionally, interferometric scattering (iSCAT) microscopy has been combined with single-particle tracking achieving nanometer precision (*18*).

Orientation-independent DIC (OI-DIC) microscopy, a method that emerged out of orientation-independent polarization microscopy (*19*), is another QPC method featuring a combination of strengths regarding its implementation, sensitivity, and resolution not present in other QPC methods. OI-DIC is an extension of the DIC concept in which the classical DIC prisms have been replaced by electronically controlled beam shearing assemblies. The beam shearing assemblies allow the DIC shear directions and biases to be switched rapidly to acquire DIC images with varying shear and bias without mechanically rotating the specimen or the prisms (*20*, *21*). OPD images are reconstructed from four to six raw DIC images without the need for phase unwrapping (*22*). The low number of required images that can be acquired at high speed make the concept compatible with live-cell imaging. OI-DIC employs partially coherent illumination with full numerical apertures (NAs) of the condenser and objective, resulting in speckle noise-free images with high phase delay sensitivity, high contrast, and good diffraction-limited resolution. Standard DIC microscopes can easily be upgraded to OI-DIC since the beam shearing assemblies fit into the DIC prism slots of conventional microscope stands. This flexibility offers a great opportunity to combine the OI-DIC with fluorescence microscopy. For example, OI-DIC has been combined with a confocal laser scanning microscope to observe chromosome condensation during mitosis (*23*). However, its performance has so far been restricted to diffraction-limited imaging in the fluorescence channel.

Here we present a correlative microscope that integrates OI-DIC into a super-resolution SMLM instrument. Our instrument is based on a commercial inverted microscope stand and achieves ∼0.05 nm OPD sensitivity with ∼2.5 nm single-molecule localization precision in the fluorescence channel. Placing the OI-DIC beam shearing assembly outside of the microscope stand enables easy integration of OI-DIC as an add-on module on most commercial inverted microscope stands compatible with standard DIC imaging. We discuss the design, characterize the two imaging modalities and demonstrate that this microscope enables 3D super-resolution fluorescence imaging in the quantitative cellular context. In addition, we show its capability of imaging specifically labeled organelles in the cellular context of living cells. Moreover, using deep-learning models to digitally stain nuclei, mitochondria, and lipid droplets in OI-DIC images of living cells, we demonstrate the potential for non-invasive continuous live-cell imaging of organelles without fluorescent labeling.

## Results

### Implementation of a correlative OI-DIC/SMLM microscope

The instrument is built based on a commercial inverted microscope stand, and a simplified schematic is shown in Fig. 1A (see Methods and fig. S1 for details). Merging of the OI-DIC and SMLM channels is realized by moving the OI-DIC recombination path outside of the microscope stand and diverting the fluorescence emission before the recombination. This ensures the OI-DIC signal passes through both beam-shearing assemblies and the fluorescence signal passes through neither (*24*). The light incidents on the channel-splitting dichroic at 8°, which gives the dichroic a notch passband at around 546 nm and similar transmission spectra for s- and p-polarized light (see fig. S1). The OI-DIC illumination is provided by a halogen lamp and an excitation filter with a passband centered at 546 nm and a bandwidth of 22 nm to match the optimal working wavelength of the beam shearing assemblies. Before acquiring raw images for OI-DIC reconstruction, the biases produced by the assemblies are calibrated as a function of voltage. Usually, 4 (4-frame mode) or 6 (6-frame mode) raw DIC images with two shear directions and multiple biases controlled by the assemblies are acquired depending on the applications. 6-frame mode is usually used in fixed cell imaging to account for unpolarized light and 4-frame mode is usually used in live-cell imaging to maximize the imaging speed (*25*) (Methods). For fluorescence imaging, four laser lines at 642 nm, 560 nm, 488 nm, and 405 nm wavelengths are coupled into the inverted microscope stand and focused into the sample plane of a 100x, 1.35 numerical aperture, silicone-oil objective. To enable 3D super-resolution imaging, a cylindrical lens is mounted on a swappable mount and near the intermediate image plane for astigmatic SMLM (*26*). Images of the OI-DIC and SMLM channels are recorded on two different sCMOS cameras, and the acquisition and analysis are automated using an open-source software package PYthon Microscopy Environment (PYME) (*27*).

**Fig. 1.**
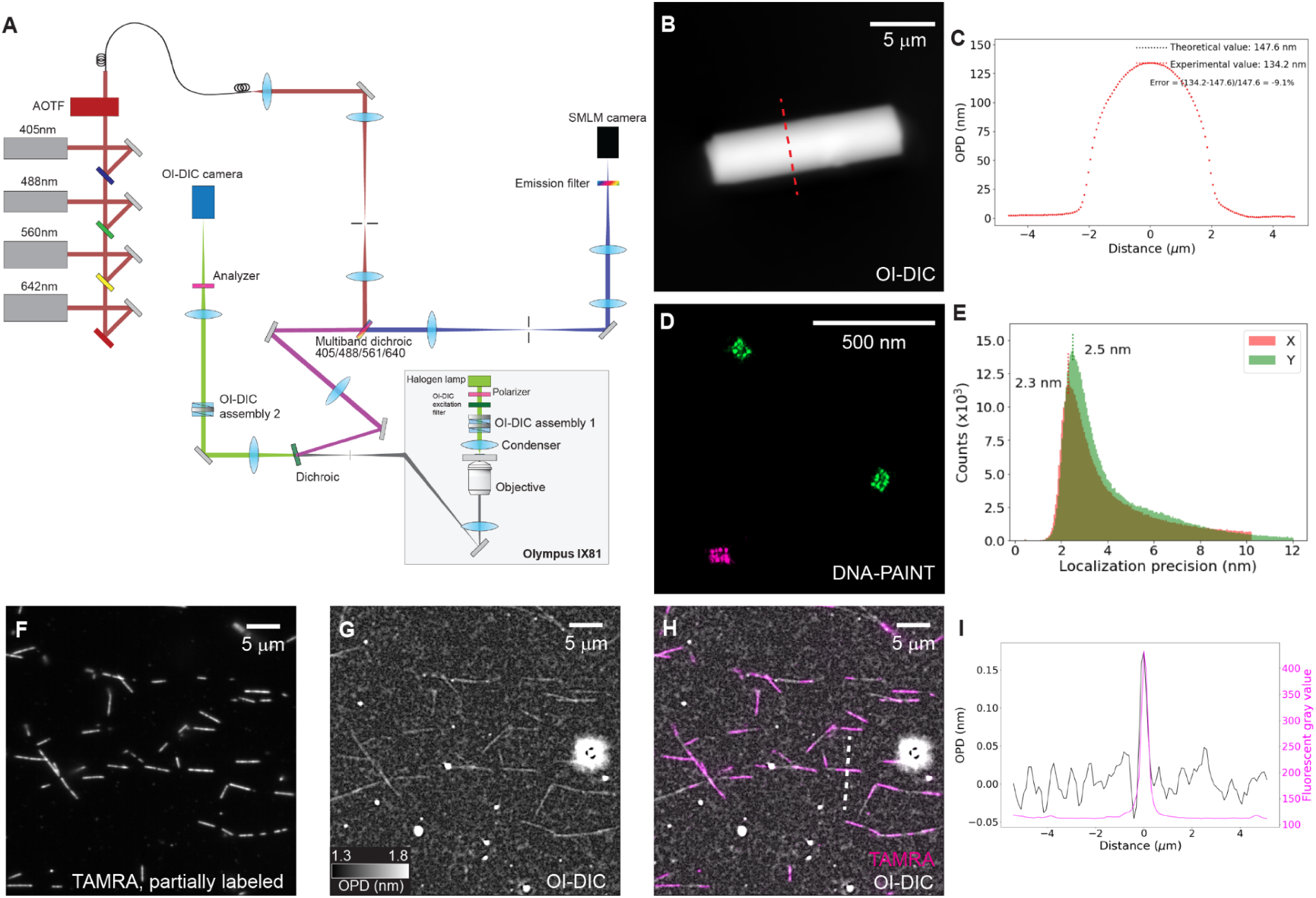
Schematic and characterization of the correlative OI-DIC/SMLM microscope. (**A**) Simplified schematic. (**B,C**) Characterization of quantitative measurement of the OI-DIC channel using a 4.1-μm-diameter glass rod. (**B**) An OI-DIC image of the 4.1-μm-diameter glass rod. (**C**) Line profile along the dashed line in **B**. (**D**,**E**) Characterization of localization precision of the SMLM channel using 20-nm-grid shaped DNA origami. (**D**) DNA-PAINT images of the DNA origami. (**E**) Histogram of localization precision of imaging the DNA origami. The lateral localization precision peaks at < 3 nm. (**F-I**) Sensitivity test using correlative fluorescence and OI-DIC imaging of in vitro microtubules. (**F-H**) Fluorescence image of discontinuously labeled in vitro microtubules (TAMRA) (**F**), OI-DIC image of the in vitro microtubules in the same field of view in **F** (**G**), overlay of **F** and **G** (**H**). (**I**) Line profile along the dashed line in **H**, indicating the visibility of the in vitro microtubules in the OI-DIC channel and the correlation with fluorescence.

Registration between the OI-DIC and SMLM channels is crucial for correlative imaging. To consider field-dependent distortions, we have implemented a shift vector map calibration by using 200-nm-diameter fluorescent beads (*28*) (see Methods and fig. S1). Although the relative positions between the two channels can drift over weeks, such drift only exists in lateral translation and the registration remains stable during the imaging acquisition. Stabilizing the imaging plane is also important for SMLM and live-cell imaging. To maintain the focal plane, a software-based drift tracking is implemented by continuously comparing the live-view and a pre-calibrated z-stack in the OI-DIC channel (see Methods and fig. S1). The lateral drift is also recorded but not maintained during the drift tracking. For SMLM imaging, the fixed cells serve as fiducials. In live-cell imaging, since the cells may move, non-fluorescent beads are added as fiducials for drift tracking. This method reduces the complexity of the hardware and can record the lateral drift for drift correction post-processing of SMLM images.

There are two major improvements over the conventional OI-DIC and fluorescence microscope. First, bringing the OI-DIC detection path outside of the microscope stand prevents interference with other modalities, making the customized OI-DIC parts a flexible add-on module for most commercial microscope stands. Second, the multicolor, high power laser excitation, swappable cylindrical lens and high-end sCMOS camera provide super-resolution applications by 3D SMLM whenever needed.

### Characterization of OI-DIC imaging and super-resolution fluorescence modes using artificial test structures

To quantitatively characterize the two imaging modalities, we imaged 4.1-μm-diameter glass rods in the OI-DIC channel (Fig. 1B,C and fig. S2) and 20-nm-grid shaped DNA origami in the SMLM channel using 2D DNA-PAINT (*29*) (Fig. 1D,E). For OI-DIC characterization, the glass rods had a refractive index of 1.560 and were embedded in the medium with a refractive index of 1.524. The intensity profile along the red dashed line matched the theoretical OPD profile with a slight deviation of 9.1% (Fig. 1C). Since this deviation was systematic and reproducible, we can safely say the OI-DIC image showed quantitative measurement of the OPD between the specimen and surrounding medium. For SMLM characterization, the 3x4 grid structures with uniform spacing of 20 nm were clearly resolvable in the DNA-PAINT image (Fig. 1D). The localization precisions peaked at 2.3 nm and 2.5 nm in x and y directions, respectively (Fig. 1E), validating the super-resolution capability of the SMLM channel.

### Testing the sensitivity of OI-DIC with *in vitro* microtubule samples

Next, we tested the sensitivity of the OI-DIC channel with discontinuously fluorescently labeled microtubules *in vitro* (Fig. 1F-I). The raw DIC images were taken by averaging 32 acquisitions for each shear and bias to reduce background noise. OI-DIC sensitivity can be defined by the smallest OPD it can detect (*30*). In the OI-DIC image, the *in vitro* microtubules were clearly visible (Fig. 1G). The OI-DIC also visualized the unlabeled segments in the absence of fluorescent labeling (Fig. 1H), suggesting this is not caused by bleedthrough between the channels (see fig. S4). This demonstrates that our OI-DIC channel has the sensitivity to detect the OPD of a single microtubule filament. Moreover, sensitivity can be quantified by the standard deviation of OPD measured when no specimen is placed on the microscope (*31*). By averaging 64 acquisitions, we find our OI-DIC has a sensitivity of detecting ∼0.05 nm OPD, being one of the most sensitive QPC microscopy (*6*) (see fig. S2).

### Correlative epifluorescence and OI-DIC imaging of cells

We then tested whether we can quantitatively image specific subcellular structures within cells using our OI-DIC. To identify the structures from the OI-DIC channel, we immunolabeled various specific organelles in fixed COS-7 cells and imaged them correlatively in the OI-DIC and epifluorescence channels. In our images, the nucleus (Fig. 2A) and mitochondria (Fig. 2B) were clearly visible in the OI-DIC channel. Microtubules (Fig. 2C) and intermediate filaments (Fig. 2D) were also visible in the OI-DIC channel with the help of a pre-extraction step performed before cell fixation (Methods). Endoplasmic reticulum (ER) was more challenging to visualize with OI-DIC (Fig. 2E) due to its small refractive index contrast compared with surroundings. The correlative imaging brings specificity to the OI-DIC signal by fluorescence and, in the opposite direction, puts specific labeled organelles in the cellular context.

**Fig. 2.**
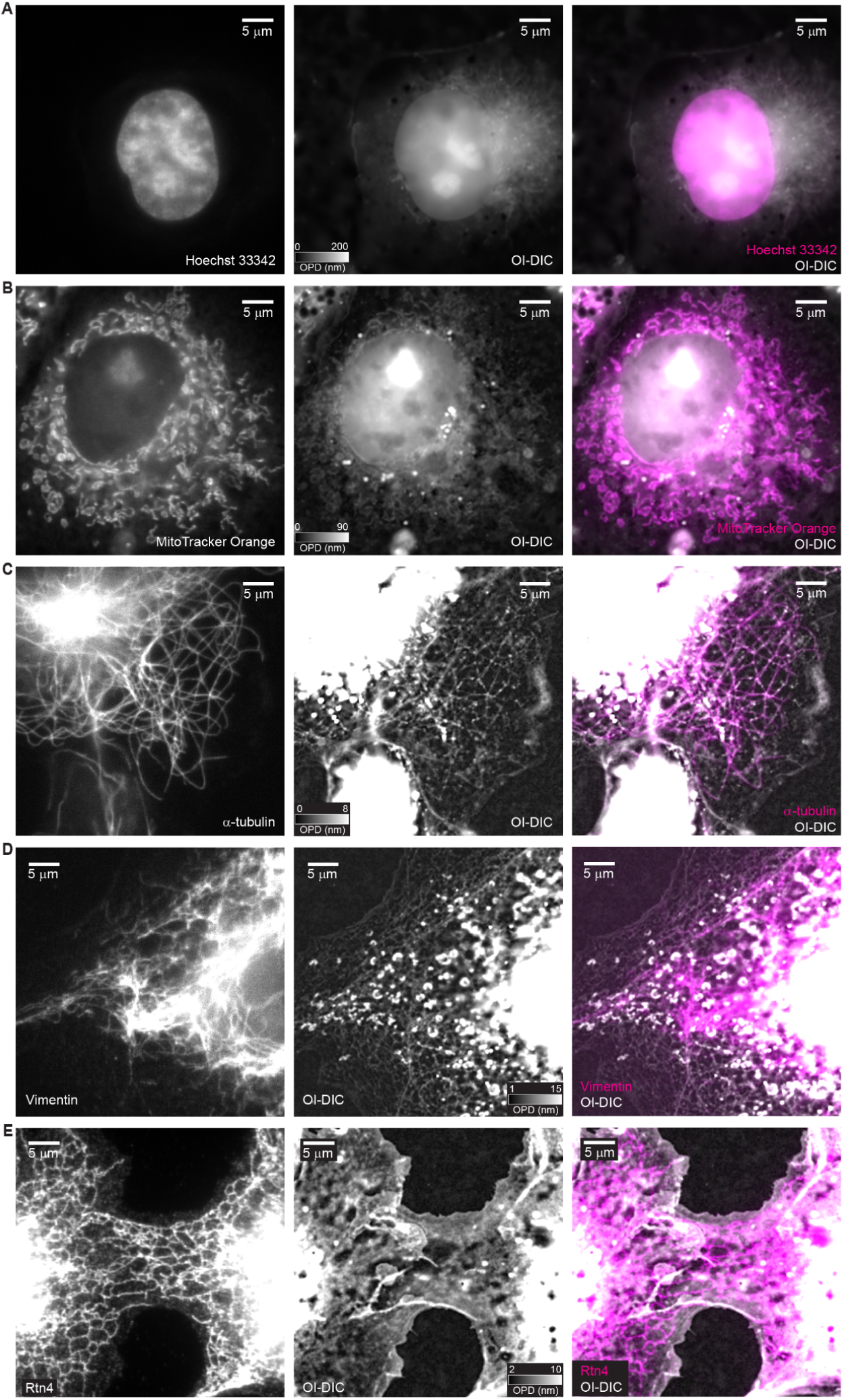
Correlative epifluorescence and OI-DIC imaging. (**A-E**) Correlative images of nucleus (**A**), mitochondria (**B**), microtubules (**C**), intermediate filaments (**D**), and ER (**E**) in fixed COS-7 cells. Columns from left to right, epifluorescence image, OI-DIC image, overlay of fluorescence and OI-DIC images.

### Correlative 3D SMLM and OI-DIC imaging in fixed cells

To test the capability of super-resolution imaging in the context of cellular structures revealed by OI-DIC, we first imaged immunolabeled microtubules in fixed COS-7 cells. Focal plane stability is crucial during SMLM imaging. To engage the software-based drift tracking, we first recorded a z-stack in the OI-DIC channel, and then the drift tracking was turned on by comparing the live-view and pre-calibrated z-stack (see Methods and fig. S1). To perform correlative imaging, we captured OI-DIC images before turning on the SMLM excitation. For SMLM imaging, we chose DNA-PAINT with fluorogenic probe Cy3B-BHQ2 due to its low background and high imaging speed (*32*). A cylindrical lens was placed near the variable square aperture to add astigmatism for 3D localization (see Methods and fig. S3). We were able to acquire a high-quality 3D super-resolution image of the microtubule network in 17 min at a fluorogenic probe concentration of 5 nM and a camera rate of 100 Hz (Fig. 3A,D). The hollow centers of microtubules can be seen in the 30-nm-thick z slice (Fig. 3D). The microtubules also were well-resolved in the OI-DIC images (Fig. B,E). Finally, the overlay was performed by applying the shift-map to the images captured from the two channels, and correlated OPD and fluorescence signals of microtubules can be seen (Fig. 3C,F).

**Fig. 3.**
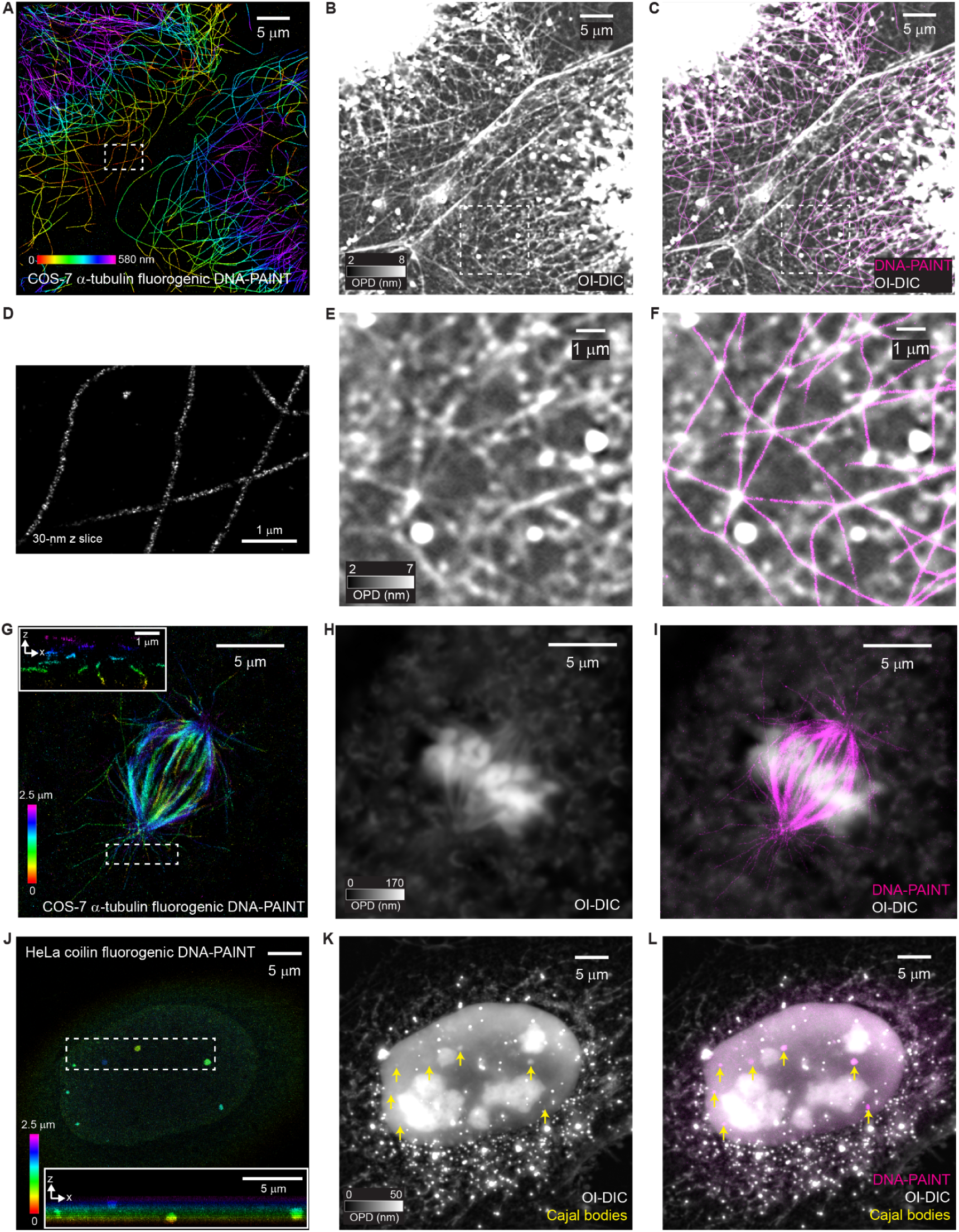
Correlative 3D SMLM and OI-DIC imaging in fixed cells. (**A**) Astigmatic 3D fluorogenic DNA-PAINT image of immunolabeled microtubules in COS-7 cells without z-stacking. (**B**) OI-DIC image of the microtubules in the same field of view in **A**. (**C**) Overlay of **A** (shown in magenta) with **B**. (**D**) Magnified view of the dashed box in **A**, representing a 30-nm-thick slice. (**E**) Magnified view of the dashed box in **B**. (**F**) Magnified view of the dashed box in **C**. (**G**) Astigmatic 3D fluorogenic DNA-PAINT image of immunolabeled spindles in a COS-7 cell with z-stacking. The inset shows the x-z cross-section of the magnified view of the dashed box. (**H**) A single slice of an OI-DIC z-stack of the spindle in the same field of view as shown in **G**. (**I**) Overlay of **G** (shown in magenta) and **H**. (**J**) Astigmatic 3D fluorogenic DNA-PAINT image of anti-coilin immunolabeled Cajal bodies in a fixed HeLa cell. The inset shows the x-z cross-section of the magnified view of the dashed box, showing the z-position of each Cajal body according to the heatmap (contrast adjusted for better visualization). (**K**) Maximum projection in the z direction of OI-DIC z-stacking images of the same field of view in **J**. Yellow arrows point to Cajal bodies. (**L**) Overlay of **J** (shown in magenta) with **K**.

We then explored whether our microscope was capable of super-resolution imaging over a thicker structure. We imaged spindles in COS-7 cells (Fig. 3G-I) and Cajal bodies in HeLa cells (Fig. 3J-L) using the same strategy, except that z-stacking was performed at a 300 nm step size (see Methods). A correction factor of 0.86 was implemented for z localizations in SMLM images to account for the refractive index mismatch between the aqueous imaging buffer and the glass-silicone oil-objective system (*26*). In the SMLM images, the astral microtubules of the spindle (Fig. 3G) can be clearly seen at different depths and the distribution of Cajal bodies in 3D can also be observed. For visualization purposes, one slice of the OI-DIC spindle image and z-direction maximum projected OI-DIC Cajal bodies z-stack were overlaid with the corresponding DNA-PAINT images, respectively. The overlay of the two channels shows good correlation between OPD and fluorescence signal, with the OI-DIC channel providing information about the cellular context.

### Continuous correlative imaging in living cells

Phototoxicity, photobleaching, imaging speed and temperature-dependent drift are major challenges for live-cell imaging. As a label-free technique, OI-DIC has inherently low phototoxicity and no photobleaching and is therefore a good candidate for imaging living cells. To demonstrate this, we imaged fluorescently labeled organelles in the cellular context by correlative OI-DIC and epifluorescence imaging in living cells. We conducted OI-DIC acquisition in 4-frame mode for faster speed and achieved around 1.1 frame per second acquisition. During acquisition, non-fluorescent beads embedded in the sample were used as fiducials for software-based drift tracking (see Methods). The OI-DIC images were captured continuously, and the fluorescence images were captured once every 5 minutes to keep phototoxicity and photobleaching at a low level (see Methods).

We first imaged actin networks in living COS-7 cells. The OI-DIC images clearly depicted the actin bundles within cells, as validated by the corresponding SiR-actin fluorescence labeling (Fig. 4A). Notably, we could capture the diverse actin dynamics in the cellular context. We examined a region near the edge of the plasma membrane and observed actin cortex redistribution along the membrane contour (Fig. 4B and Movie S1). Additionally, in another region, we observed actin extension and contraction along with filopodia protrusion and retraction (Fig. 4C and Movie S2).

**Fig. 4.**
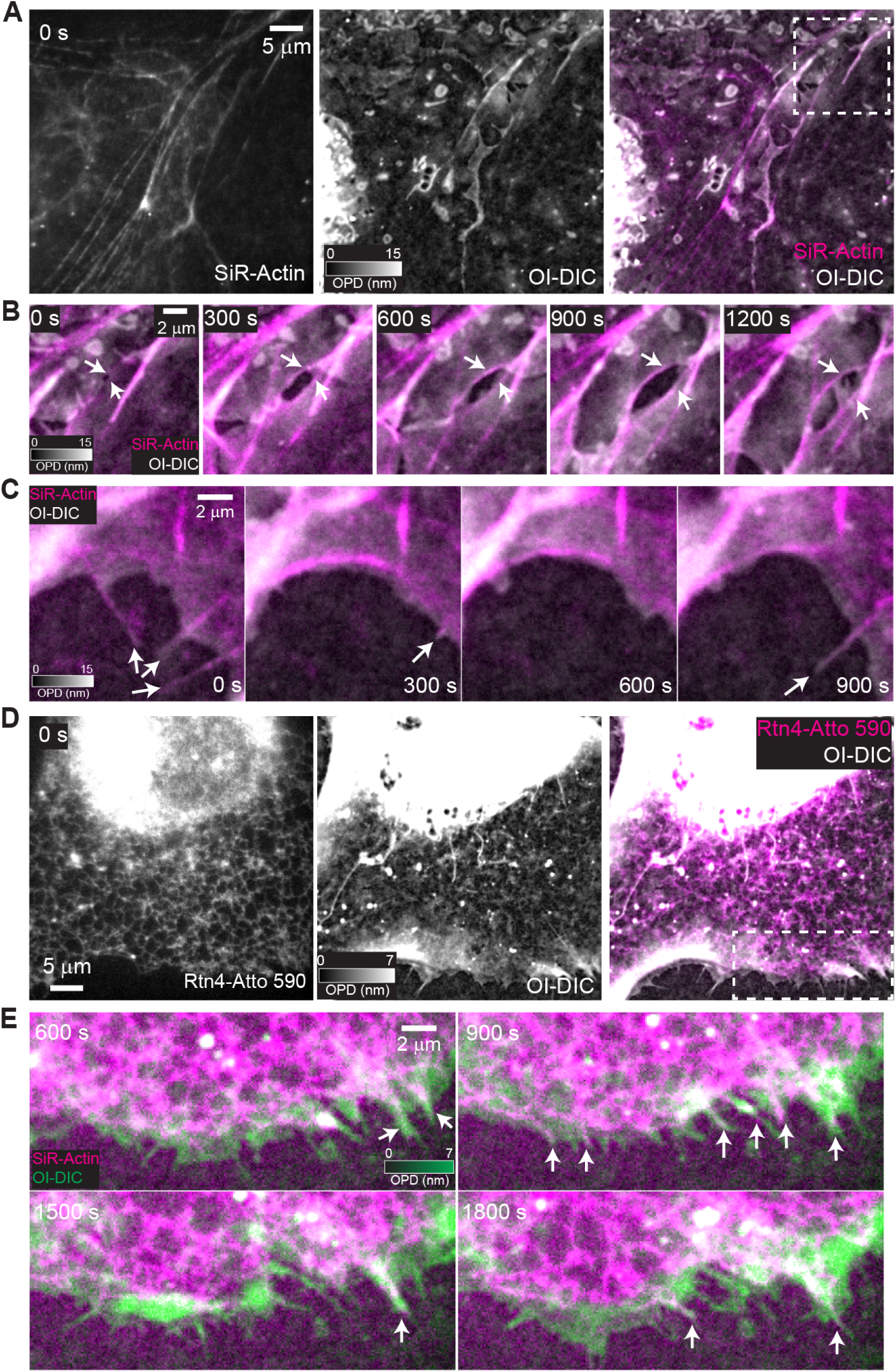
Continuous correlative imaging in living COS-7 cells. (**A**) Correlative epifluorescence image (left, SiR-Actin), OI-DIC image (middle) and overlaid image of actin networks in COS-7 cells. (**B**) Time-lapse magnified views of the dashed box in **A**, showing actin cortex redistribution along the membrane contour. (**C**) Time-lapse overlaid correlative epifluorescence and OI-DIC images, showing actin extension and contraction along with filopodia protrusion and retraction. (**D**) Correlative epifluorescence image (left, Rtn4-ATTO 590), OI-DIC image (middle) and overlaid image of ER networks in a U-2 OS cell. (**E**) Time-lapse magnified views of the dashed box in **D**, showing ER remodeling along with plasma membrane protrusions.

We next visualized the endoplasmic reticulum (ER) fluorescently labeled by Atto 590 via a ER-resident Rtn4-HaloTag construct in living gene-edited U-2 OS cells (*33*). The correlative images revealed the ER network throughout the cytoplasm of the cell (Fig. 4D) including the ER extending into dynamically remodeling filopodia (Fig. 4E and Movie S3).

### Digital staining identifies organelles from OI-DIC images

While fluorescence labeling methods provide a powerful way to highlight organelles of interest, they add to phototoxicity, are prone to photobleaching during long-term imaging, and add to the complexity of sample preparation. Artificial intelligence (AI) based ‘digital staining’ approaches are attractive alternatives to overcome these issues and have been shown to predict subcellular markers from label-free images (*34–36*). For example, fluorescent signals of various organelles have been predicted from brightfield images via different predicting models (*37*). Similarly, RI2FL predicted fluorescent labeling from live-cell 3D holotomography and has been shown to work with different cell types without model retraining (*38*). Machine learning has also been used in the work of correlative QLIPP and light-sheet fluorescence microscopy to ‘digitally stain’ nuclei and membranes for enabling segmentation and single-cell analysis (*13*).

Here we trained a Dual Squeeze-and-Excitation Residual Network (DuRDN) (*39*) to digitally stain nuclei, mitochondria, and lipid droplets in the OI-DIC data. In our experiments, we used spatially registered OI-DIC and fluorescence images of COS-7 cells for the training where the fluorescence channel served as the ground truth. Each image consisted of 416 × 416 pixels. As for the live-cell OI-DIC data above, we acquired OI-DIC data in the 4-frame mode. The raw DIC image sets, not the processed OI-DIC images, were used in the training process. We used 497 sets of images, consisting each of a fluorescence image serving as ground truth and the corresponding 4 DIC images, for training the nucleus model, 520 sets of images for training the mitochondria model, and 307 sets of images for training the lipid-droplet model (**Methods**). Testing the trained models on data sets not used in the training process showed upon visual inspection that predictions were structurally similar to the corresponding ground truth (**Fig. 5A-L**). The performance of each model was quantified by the soft Dice score (*13*) to assess the degree of overlap between digital staining and fluorescence ground truth in 47 testing image sets for the nucleus model, 48 testing image sets for the mitochondria model, and 25 testing image sets for the lipid-droplet model (**Fig. 5M; Methods**). Both the mitochondria and lipid-droplet models showed ∼0.95 soft dice scores. Such performance has been shown to be sufficiently reliable in cell phenotyping (*13*). The nucleus model showed a lower soft Dice score than the other two models. We believe the reason for this lies in the blurriness of the nuclear epifluorescence images caused by out-of-focus contributions of the bulky nuclei that are much thicker than the other two organelles. This causes a mismatch between the ‘ground-truth’ fluorescence images and the DIC images that feature inherent optical sectioning (*21*). Therefore, we only applied the mitochondria and lipid-droplet models in our subsequent studies.

**Fig. 5.**
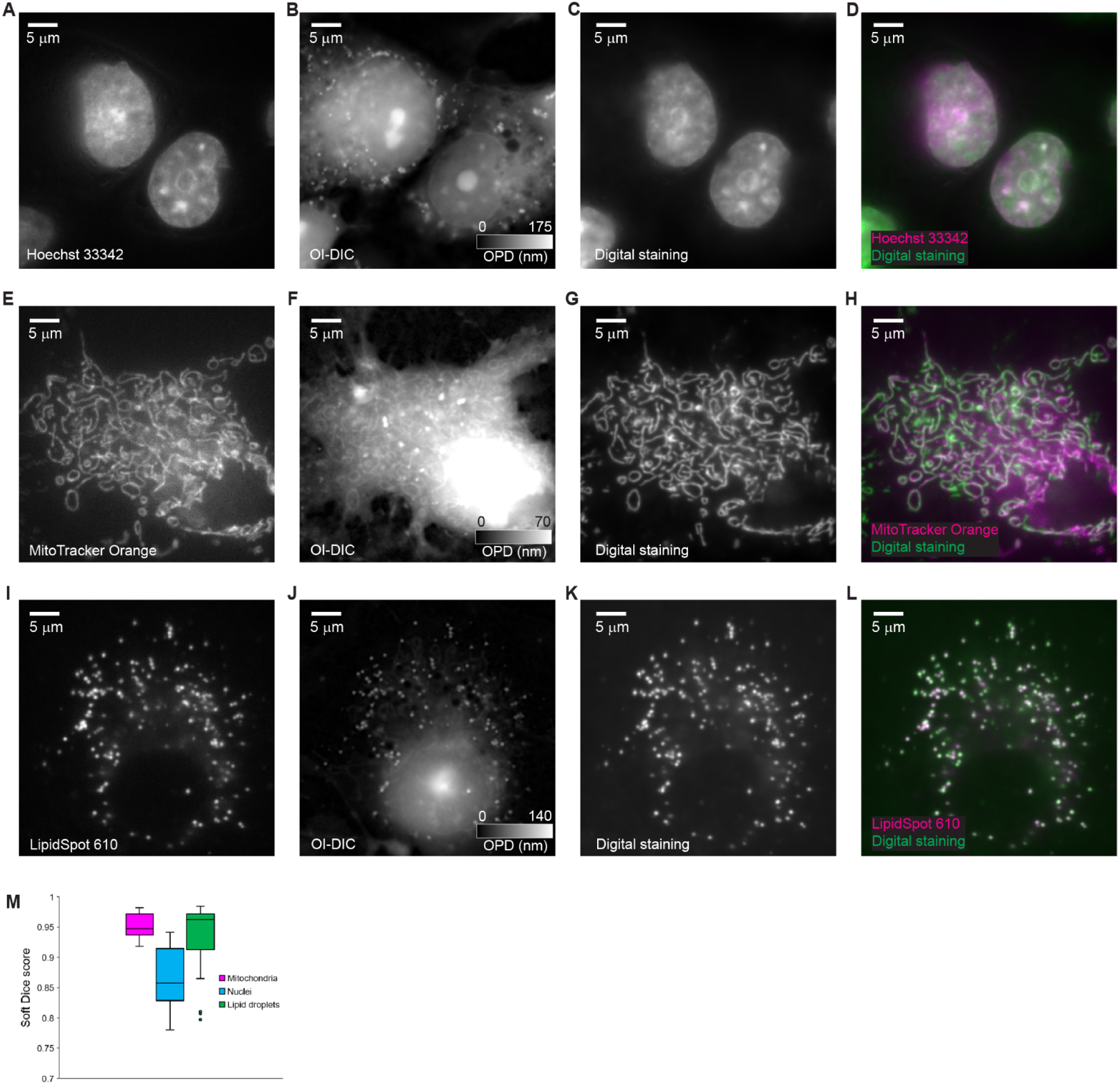
Digital staining identifies organelles from OI-DIC images. (**A-L**) Identification of nuclei (**A-D**), mitochondria (**E-H**), and lipid droplets (**I-L**) in COS-7 cells. (**A, E, I**) Epi-fluorescence images of nuclei (Hoechst), mitochondria (MitoTracker Orange), and lipid droplets (LipidSpot 610) in COS-7 cells, respectively, serving as ground truth. (**B, F, J**) OI-DIC images of the fields of view shown in **A**, **E**, **I**, respectively. (**C, G, K**) Digitally stained images generated from **B**, **F**, **J** using the nucleus, mitochondria, and lipid-droplet deep learning models, respectively. (**D, H, L**) Overlay of digital staining and fluorescence ground truth images, showing that the deep learning predictions were structurally similar to the corresponding ground truth. (**M**) Digital staining performance quantified by the soft Dice score.

### Mitochondria and lipid-droplet dynamics in living cells observed by digital staining

Using digital staining, OI-DIC enables continuous live-cell imaging of organelle dynamics with minimal phototoxicity. To demonstrate this, we continuously imaged living COS-7 cells in the OI-DIC channel at 1-s time resolution for ∼17 minutes, resulting in 1,003 time points. We then applied our pre-trained mitochondria and lipid-droplet models to the images, generating 3-channel data that way (**Fig. 6A-E**). The line profile in the magnified region of the dashed box in **Fig. 6A** shows good correlation between the OI-DIC signal and digital stainings (**Fig. 6B**). In this line profile, the OPD curve exhibits four peaks, with the first peak correlating with the peak of the lipid-droplet digital staining curve and the second, third, and fourth peak correlating with the three peaks of the mitochondria digital staining curve (**Fig. 6B**). As a control, mitochondria were stained with MitoTracker Orange CMTMRos and additionally imaged in the fluorescence channel once every 5 minutes. The visual similarity of digital staining and fluorescence validated the mitochondria prediction and confirmed our previous observation (**Fig. 6A**). In addition, the live-cell images clearly captured mitochondria and lipid-droplet dynamics (**Fig. 6A-E** and **Movie S4, S5**). For example, we observed a mitochondrion reorganizing from a rod shape to a triangular disk shape and back to a rod shape (**Fig. 6C, D**). Moreover, we could visualize mitochondria fission and fusion processes (**Fig. 6C, E**).

**Fig. 6.**
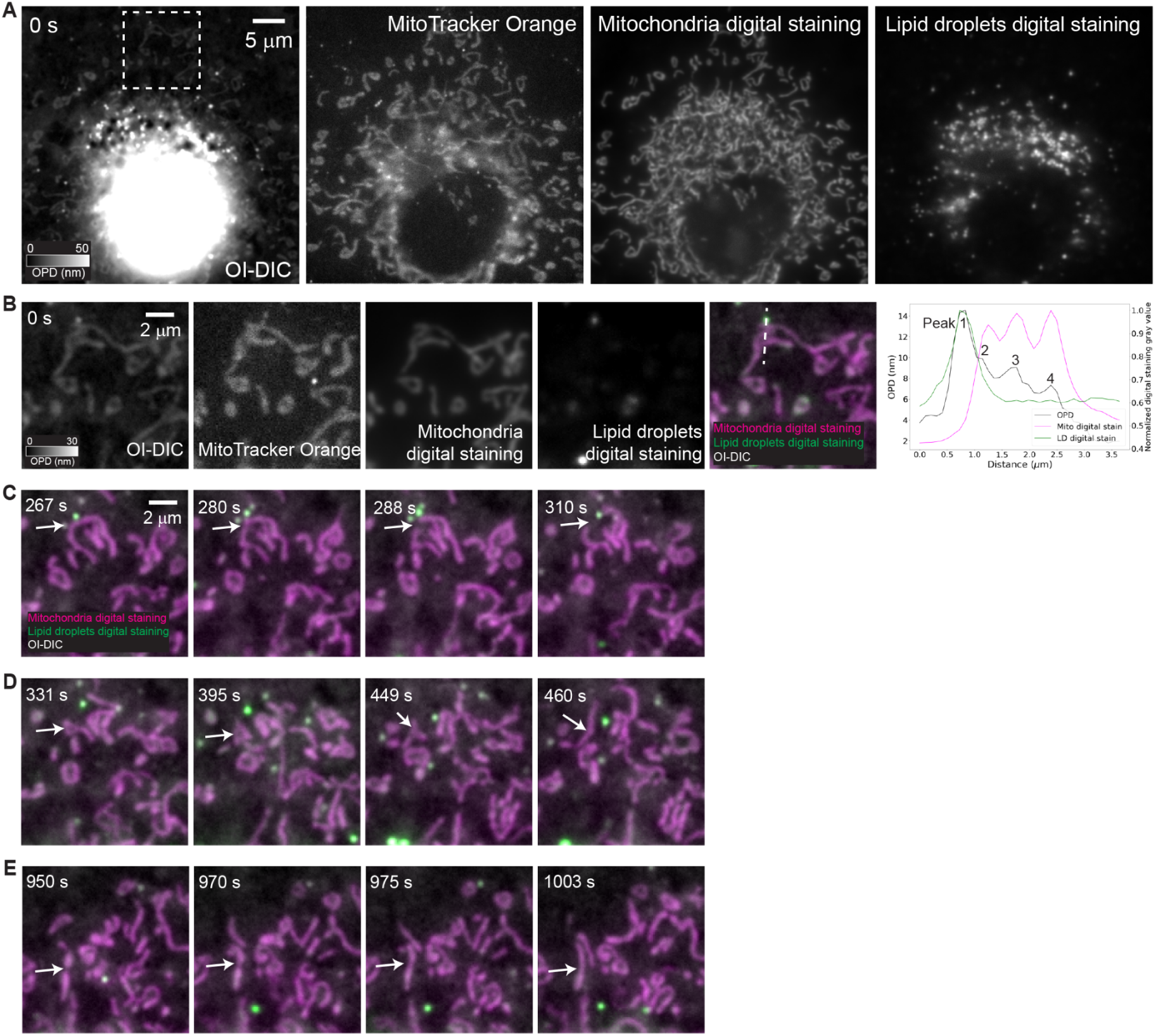
Mitochondria and lipid-droplet dynamics in living cells observed by digital staining. (**A**) Correlative epifluorescence and OI-DIC images of mitochondria and digital staining of mitochondria and lipid droplets in a COS-7 cell. (**B**) Magnified view of the dashed box in **A**. The line profile shows good correlation between OI-DIC and digital stainings. (**C-E**) Time-lapse magnified view of the dashed box in **A**, showing mitochondria fission (**C**), transition from a rod shape to a triangular disk shape and back to a rod shape (**D**), and fusion (**E**).

## Discussion

We have shown that our implementation of correlative OI-DIC/SMLM enables 3D super-resolution imaging of specifically labeled molecules in the quantitative cellular context with an OPD sensitivity of ∼0.05 nm and a single-molecule localization precision of ∼2.5 nm. The obtained OPD sensitivity compares favorably to the reported values of other QPC methods: digital holographic microscopy (DHM), based on a backscattering geometry, has demonstrated 0.9 nm OPD sensitivity (*40*, *41*); Quadriwave lateral shearing interferometry (QWLSI) has demonstrated sub-1nm OPD sensitivity (*42*). We have demonstrated the application of imaging both fixed and living cells and shown continuous live-cell imaging with mitochondria and lipid droplets digital staining by deep learning. The image acquisition and analysis have been implemented in PYME and are ready to be used for biological applications.

However, it is worth pointing out that we have not yet achieved the performance limit of this microscope. The OI-DIC channel could be further improved by optimizing the liquid crystal of a faster settling speed at a longer wavelength for 3D live-cell imaging with lower photodamage and optimizing printed nanostructured DIC prisms (*43*, *44*). We also expect advances in theoretical studies of OI-DIC to decouple 3D refractive index maps based on 3D partially coherent point spread function (PSF) (*45*) from the OPD measurements, which is the integral of the refractive index of the specimen along the axial direction. With OI-DIC’s inherent characteristics of good resolution and optical sectioning, this may enable software-based tomographic imaging with a simple OI-DIC module. The fluorescence channel could also be further improved by implementing simultaneous multicolor imaging by adding image splitters before the camera to place up to three detection channels on the camera chip. Moreover, deep learning could be applied to identifying more organelles such as microtubules and ER, enabling simultaneous observation of multiple targets. Additionally, deep learning has the potential to trigger live-cell fluorescence acquisitions in response to specific cellular events. Such a function may enhance the recently developed ‘smart microscope’ involving streaming analysis and event-driven triggering (*46*, *47*).

Our development represents an important step for correlative QPC and super-resolution fluorescence imaging. With the advances contained in this work and improvements mentioned above, our approach opens the way for biological applications that we have not yet explored in this study. A recent study presents unclearing microscopy, a new type of expansion microscopy (ExM) with absorptive or phase contrast in transmitted light microscopy (*48*). OI-DIC has great potential to interpret the 3D composition of unclearing microscopy samples. Similarly, studies such as phase separation and particle tracking in the context of surrounding cellular structures in the living cells could also be conducted with this correlative microscope. Additionally, the OI-DIC can be implemented as an add-on module at the side port of most inverted microscopes to achieve correlative imaging. Such imaging of, for example, signaling dynamics and cell differentiation, is valuable for addressing open questions in developmental and cell biology (*13*).

## MATERIALS AND METHODS

### Microscope setup

The instrument is set up based on an inverted microscope stand (Olympus, IX81) using a silicon-oil immersion objective (Olympus, UPlanSApo 100x/1.35 NA) . The microscope is equipped with a piezo objective actuator (Physik Instrumente, P-721.CLQ) for z-focusing and a motorized stage (Marzhauser Wetzlar, SCAN IM) for xy-positioning of the sample. For live-cell imaging, a stage-top incubator (Tokai Hit, STXF-WSKMX-SET) is placed on the motorized sample stage for environmental control. The OI-DIC and SMLM channels are combined outside of the microscope stand using a dichroic mirror (Semrock, Di03-R685-t1-25x36) (see fig. S1). Two beam shearing assemblies are custom-built for the OI-DIC channel and matched to the dimension of Olympus DIC prisms (*25*). The condenser-side assembly consists of two Wollaston prisms and two liquid crystals (ARCoptix, Neuchatel, Switzerland), with a liquid crystal retarder adjusting biases and a liquid crystal rotator adjusting polarization directions. The objective-side assembly is placed outside of the microscope, between the dichroic mirror and the OI-DIC camera, and consists of two Wollaston prisms and a liquid crystal rotator. For OI-DIC imaging, the sample is illuminated with a halogen lamp (Olympus, U-LH100L-3) mounted on the illumination pillar (Olympus, IX2-ILL100) with an excitation bandpass filter centered at 546 nm (Chroma, ET546/22x). Before reaching the sample, the illumination beam passes through a linear polarizer (Thorlabs, LPVISE100-A), a quarter-wave plate (Olympus, WI-TP137), the internal OI-DIC beam shearing assembly described above (*25*), and a dry condenser (Olympus, U-TLD). The light collected by the objective exits the microscope stand through the tube lens and side port, passes through the dichroic mirror, and is then re-imaged to the infinity space by an additional tube lens (Olympus, U-TLU). The external OI-DIC beam shearing assembly is placed near the image of the back focal plane of the objective matching the interference plane of the DIC prism by moving the assembly along the beam axis. The final image is formed by another tube lens (Olympus, U-TLU) and an analyzer (Thorlabs, LPVISE100-A) onto an sCMOS camera (Excelitas, pco.edge 4.2 LT).

For fluorescence imaging, four laser lines at wavelengths of 642 nm (MPB Communications), 560 nm (MPB Communications), 488 nm (Coherent OBIS 488, 150 mW) and 405 nm (Coherent OBIS 405, 100 mW) pass through an acousto-optical tunable filter (AOTF; AA Optics) used to adjust the laser intensities. The laser output of the AOTF is coupled into a square-core multimode fiber (Thorlabs, M103L05), which is vibrated to average out speckle patterns and achieve a uniform intensity at the fiber output. The fiber output is then imaged into the focal plane of the objective. The fluorescence emission is collected by the objective, reflected by the dichroic mirror, and imaged onto another camera (Hamamatsu, ORCA-Flash 4.0 V3). A multiband dichroic mirror (Chroma, ZT405/488/561/640) separates the fluorescence excitation from the emission. Detection spectra can be further selected by adding different emission filters in front of the camera. A variable square aperture (Owis, SP40) is placed at the intermediate image plane to adjust the size of the detected field of view. For 3D SMLM measurements, a cylindrical lens (Thorlabs, f = 1000 mm) is placed near the intermediate image plane. The system is controlled by the open-source software package PYME (*27*).

### Shift-map calibration

The calibration was implemented by using 200-nm-diameter fluorescent beads (Thermo Fisher Scientific, F8807) on a coverslip. The fluorescence and OI-DIC raw images were simultaneously recorded and the position difference of each bead between the two channels was measured (see fig. S1). Using smoothing spline interpolation, a shift vector map was created. In the fitting process, the fluorescence channel was set as the reference channel.

### Focus lock

A software-based drift tracking method, implemented in PYME, was used to stabilize the focal plane (see fig. S1). Before imaging, a z-stack was taken in the label-free channel at a step size of 200 nm. During the experiment, transmitted-light images were acquired with the OI-DIC camera for drift tracking. These images were cross-correlated with the initial z-stack to calculate the lateral and axial offsets. The lateral offset was recorded for later correction in post-processing and the axial offset was corrected by the piezo objective actuator.

### Sample preparation

Microscope characterization sample preparation.

#### Glass rod

Short segments of glass rods, normally used as spacers in liquid-crystal cells, were embedded in Fisher Permount mounting medium (Fisher Scientific). The refractive indices of the glass rods and the Permount mounting medium were measured at wavelength 546 nm with a Jamin–Lebedeff microscope (Zeiss, Germany), and were determined to be 1.560 and 1.524, respectively. A drop of the glass rods suspended in the mounting medium was placed between a microscope slide and a 0.17-mm thick coverslip.

#### DNA origami

Buffer A+ (10 mM Tris pH 8, 100 mM NaCl and 0.05% Tween-20) and buffer B+ (10 mM MgCl2, 5 mM Tris-HCl pH 8, 1 mM EDTA and 0.05% Tween-20) were used to prepare the DNA origami sample. When preparing the sample, a square coverslip (18x18 mm No. 1.5H) and glass slide (3”x1”x1mm) were sandwiched together with two strips of double-sided tape to form a flow chamber with an inner volume of ∼20 μl. First, 20 μl of biotin-labeled BSA (1 mg/mL in buffer A+) was flushed into the chamber and incubated for 2 min. Then, the chamber was washed with 40 μl of buffer A+ and 20 μl of streptavidin (0.1 mg/mL in buffer A+) was flushed in and incubated for 2 min. After washing with 40 μL of buffer A+ then 40 μl of buffer B+, 20 μl of biotin-labeled DNA origami (1:100 dilution in buffer B+) was flushed into the chamber and incubated for 2 min. Finally, the chamber was washed with 40 μL of buffer B+ and 20 μL of the imager solution at a concentration of 2.5 nM was flushed in. The chamber was sealed with two-component silica prior to imaging. The DNA origami sample preparation protocol was previously reported in (*29*).

#### *In vitro* microtubules, fully labeled, GMPCPP-stabilized microtubules

Biotinylated and TAMRA-labeled microtubules were polymerized by incubating a solution containing 1 mM guanylyl 5’-α,β-methylenediphosphonate (GMPCPP), 1 mM MgCl₂, 8 μM unlabeled bovine tubulin, 1 μM biotinylated tubulin, and 1 μM TAMRA-labeled tubulin in BRB80 buffer. The solution was first incubated on ice for 5 min, followed by polymerization at 37 °C for 30 min. Polymerization was stopped by adding 100 μL of room-temperature BRB80. To remove unpolymerized tubulin, microtubules were pelleted by ultracentrifugation at 126,000 × g for 5 min at room temperature. The supernatant was carefully removed, and the pellet was resuspended in 200 μL of room-temperature BRB80 by gentle pipetting to prevent microtubule shearing.

#### *In vitro* microtubules, discontinuously labeled with unlabeled extensions

The pre-formed fluorescent seeds as described above were used to nucleate unlabeled tubulin extensions. The seed solution was added to a polymerization mixture containing 4 μM unlabeled bovine tubulin, 1 mM GTP, 5 mM dithiothreitol (DTT), and 0.5 μM biotinylated tubulin in the BRB80 buffer. This mixture was incubated at 37 °C for 20 min to promote growth of the unlabeled extensions. The reaction was then stabilized by adding paclitaxel (taxol) to a final concentration of 1 μM. Finally, the microtubules were pelleted by ultracentrifugation (126,000 × g, 5 min, room temperature) to remove unincorporated tubulin and resuspended in BRB80 solution containing 1 μM taxol.

#### *In vitro* microtubules, microscopy sample preparation

For both assays, microtubules were immobilized on antibiotin antibody-coated silanized coverslips, prepared as previously described (*49*). The presence of surface-bound microtubules was confirmed via simultaneous interference reflection microscopy (IRM) and total internal reflection fluorescence (TIRF) microscopy (*50*) prior to OI-DIC measurements.

### Cell sample preparation

#### Cell culture

COS-7 cells (ATCC, CRL-1651, batch #63624240) were grown in DMEM (Gibco, 21063029) supplemented with 10% fetal bovine serum (FBS; Gibco, 10438026) at 37 °C with 5% CO_2_. The U-2 OS Rtn4-Halo CRISPR cell line was developed as previously reported (*33*) and grown in McCoy’s 5A (Gibco, 16600082) supplemented with 10% FBS at 37 °C with 5% CO_2_.

#### Coverslip and MatTek dish cleaning

No. 1.5H, 18-mm-diameter round coverslips were used for fixed-cell sample preparation and MatTek glass bottom dishes (MatTek Life Sciences, P35GC-1.5-20-C and P35GC-1.5-14-C) were used for live-cell sample preparation. Both were cleaned in a sonic bath immersed in 1 M KOH for 15 min and then rinsed three times with Milli-Q water. They were then sterilized with 70% ethanol followed by poly-L-lysine (Sigma-Aldrich, P4707) coating for 10 min. After that, coverslips were rinsed with sterilized PBS (1xPBS; Gibco, 10010023) before adding media and cells, while MatTek dishes were further coated with 2-μm-diameter beads (Thermo Fisher Scientific, C37248) in PBS at a dilution of 1:500 for 10 min and then rinsed with PBS before adding media and cells.

#### Nucleus labeling

COS-7 cells were grown on pre-cleaned and poly-L-lysine (Sigma-Aldrich, P4707) coated coverslips and live-labeled with Hoechst 33342 (Tocris Bioscience, 5117) diluted in warm media for 10 min at 37 °C with 5% CO_2_. Cells were then rinsed three times with warm media and fixed with 3% paraformaldehyde (PFA; Electron Microscopy Sciences, 15710) and 0.1% glutaraldehyde (GA; Electron Microscopy Sciences, 16019) for 15 min at room temperature. Lastly, the samples were rinsed with PBS three times and stored in PBS at 4 °C until imaged.

#### Mitochondria labeling

For fixed cell samples, COS-7 cells were grown on pre-cleaned and coated coverslips and live labeled with 500 nM MitoTracker Orange CMTMRos (Thermo Fisher Scientific, M7510) diluted in warm media for 30 min at 37 °C with 5% CO_2_. Cells were then rinsed three times with warm media and fixed with 3% PFA + 0.1% GA for 15 min at room temperature. Lastly, the samples were rinsed with PBS three times and stored in PBS at 4 °C until imaged. For live-cell samples, COS-7 cells were grown on pre-cleaned and coated MatTek dishes and live labeled with 200 nM MitoTracker Green FM (Thermo Fisher Scientific, M7514) diluted in warm media for 45 min at 37 °C with 5% CO_2_. Cells were then rinsed three times with warm media. The media was replaced with Live Cell Imaging Solution (Gibco, A59688DJ) supplemented with 3% FBS. Cells were then imaged immediately while being maintained at 37°C.

#### Microtubules labeling

For DNA-PAINT imaging, COS-7 cells were grown on pre-cleaned and coated coverslips and pre-extracted with prewarmed (37 °C) 0.05% saponin diluted in cytoskeletal buffer (CBS; 10 mM MES, 138 mM KCl, 3 mM MgCl_2_, 2 mM EGTA, 320 mM sucrose) for 45 s. The pre-extraction buffer was then removed. The cells were then fixed with prewarmed (37 °C) 3% PFA + 0.1% GA in CBS buffer for 15 min, rinsed with PBS three times, incubated in cytoskeletal block buffer (PBS + 3% BSA + 0.2% TX-100) for 30 min, and incubated with mouse anti-α-tubulin primary antibody (Sigma-Aldrich, T5168) at a dilution of 1:1,000 in antibody-dilution buffer (PBS + 1% BSA + 0.2% TX-100) overnight at 4 °C. The cells were then washed three times for 5 min each with cytoskeletal wash buffer (PBS + 0.05% TX-100) and incubated with oligonucleotide-conjugated goat anti-mouse IgG secondary antibody (Jackson ImmunoResearch, 111-005-144) at a dilution of 1:200 in antibody-dilution buffer for 1 h at room temperature. The secondary antibody was conjugated to oligonucleotide docking strands using azide-DBCO click chemistry (*29*). Lastly, the cells were washed three times for 5 min each with cytoskeletal wash buffer, post-fixed with 3% PFA + 0.1% GA in CBS for 10 min, rinsed with PBS three times and stored in PBS at 4 °C until imaged. For fixed cell epifluorescence imaging, same protocol was used except the cells were incubated with goat anti-mouse-IgG-ATTO 647N (Sigma-Aldrich, 50185) for the secondary antibody incubation. For live-cell imaging, COS-7 cells were grown on pre-cleaned and coated MatTek dishes and live labeled with 1 μM SiR-Tubulin with 10 μM verapamil (Cytoskeleton, CY-SC002) in warm media for 1 h at 37 °C with 5% CO_2_. The cells were then rinsed once with warm media. The media was then replaced with Live Cell Imaging Solution supplemented with 3% FBS. Lastly, the cells were imaged immediately while being maintained at 37 °C.

#### Cajal bodies labeling

HeLa Kyoto cells (from ATCC) were cultured in DMEM GlutaMAX (Gibco) supplemented with 10% FBS (Gibco) and 1% PenStrep (Gibco) at 37 °C with 5% CO_2_. 0.2 million cells were seeded onto 12-well plates (Falcon) on 18 mm, #1.5 round coverslips (Neuvitro) and grown for 24 h. Cells were then fixed in 4% PFA in PBS for 10 min, permeabilized and blocked by blocking buffer (0.1% Triton X-100 and 3% BSA in PBS) for 15 min, incubated with Coilin antibody (1:1000 dilution, Abcam, ab87913) in blocking buffer for 1 h, washed, and stained with oligonucleotide-conjugated goat anti-mouse IgG secondary antibody (Jackson ImmunoResearch, 111-005-144) (1:1000 dilution) for 1 h in blocking buffer. The stained cells were fixed again in 4% PFA in PBS for 10 min, and kept at 4 °C before imaging.

#### Intermediate filaments labeling

Same protocol as fixed cell microtubules labeling was used except the cells were incubated with rabbit anti-vimentin (ProteinTech, 10366-1-AP) for the primary antibody incubation and goat anti-rabbit-IgG-ATTO 647N (Rockland Immunochemicals, 611-156-122) for the secondary antibody incubation.

#### ER labeling

For fixed cell samples, COS-7 cells were grown on pre-cleaned and coated coverslips and fixed with 3% PFA + 0.1% GA for 15 min at room temperature. The cells were then rinsed with PBS three times, incubated in permeabilization buffer (PBS + 0.5% TX-100) for 3 min, incubated in block buffer (PBS + 0.1% TX-100 + 1% BSA) for 60 min, and incubated with rabbit anti-Rtn4 primary antibody (Abcam, ab47085) at a dilution of 1:200 in block buffer overnight at 4 °C. The cells were then washed three times for 5 min each with 1xPBST (PBST) and incubated with goat anti-rabbit-IgG-ATTO 594 secondary antibody (Sigma-Aldrich, 77671) at a dilution of 1:200 in block buffer for 1 h at room temperature. Lastly, the cells were washed three times for 5 min each with PBST, post-fixed with 3% PFA + 0.1% GA for 10 min, rinsed with PBS three times and stored in PBS at 4 °C until imaged. For live-cell samples, channel slides (ibidi, 80187) were coated with 2-μm-diameter beads (Thermo Fisher Scientific, C37248) in gelatin (Sigma-Aldrich, G1890-100G) at a dilution of 1:1,000 for 1 h at 37 °C. U-2 OS Rtn4-Halo cells were then grown on the channel slides and live labeled with 1 μM ATTO 590-chloroalkane diluted in warm media for 1 h at 37 °C with 5% CO_2_. The ATTO 590-chloroalkane probe was developed as previously reported (*33*). Cells were then rinsed three times with warm media and allowed to recover for 1 h at 37 °C with 5% CO_2_. The media was then replaced with Live Cell Imaging Solution supplemented with 3% FBS. Lastly, the cells were imaged immediately while being maintained at 37 °C.

#### Lysosome labeling

Same protocol as live-cell microtubules labeling was used except using 1 μM SiR-Lysosome with 10 μM verapamil (Cytoskeleton, CY-SC012) as staining solution.

#### Actin labeling

Same protocol as live-cell microtubules labeling was used except using 1 μM SiR-Actin with 10 μM verapamil (Cytoskeleton, CY-SC001) as staining solution.

#### DNA-PAINT buffer

For fixed cell samples, a DNA-PAINT buffer (PBS + 500 mM NaCl + 20 mM Na_2_SO_3_ + 1 mM Trolox) was freshly made. Trolox (Sigma-Aldrich, 238813) aliquots were stored at -20 °C at a concentration of 50 mM in DMSO (Thermo Fisher Scientific, D12345). Before imaging, the cells were rinsed with DNA-PAINT buffer three times, and fluorogenic imager probes Cy3B-BHQ2 were added to the cell at a proper dilution in the buffer (5 nM final concentration for imaging microtubules without z-stacking, 4 nM final concentration for imaging spindles with z-stacking, and 50 nM final concentration for imaging Cajal bodies). To calibrate experimental PSF, the 100-nm-diameter beads (Thermo Fisher Scientific, F8801) were added to the sample at a dilution of 1:1,000,000. The buffer composition and imager probes were previously reported in (*32*).

### Deep learning / digital staining

Four spatially registered, raw DIC and fluorescence images of living COS-7 cells were used for the training. Each image consisted of 416 × 416 pixels. A total of 1444 image sets were collected and utilized in this work, covering nuclei, mitochondria, and lipid droplets. Specifically, 544 nucleus image sets were included–497 for training and 47 for testing. Similarly, 568 mitochondria image sets were collected, with 520 allocated for training and 48 for testing. Lastly, 332 lipid droplet image sets were incorporated, all 307 used for training and 25 reserved for testing. Separate neural networks are trained for each individual organelle category. A convolutional neural network architecture, termed as Dual Squeeze-and-Excitation Residual Network (DuRDN), was employed in this work for automatic staining of nuclei, mitochondria, and lipid droplets in the collected OI-DIC data. DuRDN (*39*) employs a conventional U-Net backbone. It includes an encoder for extracting image features, a decoder for reconstructing image details, and skip connections between corresponding layers to preserve spatial information that might be lost during downsampling. Additionally, residual dense connections (RDC) and squeeze-excitation attention (SEA) mechanisms make it powerful to extract the very detailed information of the OI-DIC data. RDC aggregates the input features of previous concatenated convolutional layers, thus enabling enhanced image feature reuse and information extraction. SEA can recalibrate image feature weights across both channel-wise and spatial dimensions, therefore enhancing the model’s focus in critical regions of interests (ROI). All these designs improve the image processing ability of DuRDN and make it a more powerful tool over the standard U-Net architecture.

### Image acquisition

#### OI-DIC acquisition

Kohler illumination was adjusted and the waiting time for the liquid crystal settling was set to 100 ms. Before acquiring raw images, the delay produced by the liquid crystal retarder was calibrated as a function of voltage. For fixed cell imaging, 6 raw DIC images of two orthogonal shear directions and -0.15π, 0, 0.15π biases were acquired (6-frame mode), and for live-cell imaging, 4 raw DIC images of two orthogonal shear directions and -0.3π, 0.3π biases were acquired to increase the acquisition speed (4-frame mode). Several raw frames of the same shear direction and bias could be captured for averaging to reduce the background noise whenever necessary (frame averaging). A background OI-DIC image was taken in the end with the same settings for reconstruction by defocusing the target or by moving the field of view to an empty area in the sample.

#### DNA-PAINT image acquisition for DNA origami

The imaging was performed at a camera exposure time of 150 ms and 560-nm laser illumination with an intensity of 1 kW cm^-2^ delivered to the sample. 25,000 frames were recorded.

#### Correlative OI-DIC and epifluorescence acquisition for fixed cells

The OI-DIC and fluorescence acquisitions were initiated simultaneously controlled by a self-programed acquisition protocol in PYME. OI-DIC images were acquired with a camera exposure time of 25 ms in 6-frame mode using 4-frame averaging. Epifluorescence imaging was performed at 100 ms camera exposure time. Lastly, the OI-DIC background image was acquired.

#### Correlative OI-DIC and DNA-PAINT acquisition for fixed cells

A cylindrical lens (f = 1000 mm) was added to the fluorescence emission path to introduce astigmatism. An experimental PSF was calibrated first by taking a z-stack of 100-nm-diameter beads in the sample with 50 nm step size and 150 slices. After that, the OI-DIC image was acquired with a camera exposure time of 25 ms in 6-frame using 4-frame averaging, and then the software-based drift tracking was engaged. DNA-PAINT imaging was performed at 100 Hz and 560-nm laser illumination with an intensity of 13 kW cm^-2^ delivered to the sample. Lastly, the OI-DIC background reference image was acquired.

#### Correlative OI-DIC and epifluorescence acquisition for living cells

The software-based drift tracking was engaged before data acquisition started. OI-DIC and fluorescence data acquisition was controlled by an acquisition protocol in PYME. The OI-DIC images were acquired continuously at 4-frame mode and 1-frame averaging with a camera exposure time of 25 ms. Epifluorescence excitation and acquisition were turned on around every 5 min with the same settings as the epifluorescence imaging mentioned above.

### Image processing

Image data were recorded and analyzed with a custom-programmed open-source software package PYthon Microscopy Environment (PYME) (*27*). For two-channel overlaying, OI-DIC images were rolling-ball background subtracted using built-in functions in Fiji (*51*) before overlaying. For 3D DNA-PAINT images, localizations were performed based on experimental PSF (*52*) and were then drift-corrected first based on recorded lateral drift by software-based drift tracking and then by a redundant cross-correlation (*53*). For deep learning training and digital staining, the network was run on the Yale High Performance Computing cluster to get the final result.

## Supplementary Materials

Supplementary Fig. S1. Optical setup and characterization of the correlative microscope.

Supplementary Fig. S2. Characterization of the OI-DIC channel.

Supplementary Fig. S3. PSF calibration of the fluorescence channel.

Supplementary Fig. S4. OI-DIC imaging of unlabeled COS-7 cells with z-stacking.

Supplementary Movie S1. A live-cell movie of actin in a COS-7 cell.

Supplementary Movie S2. Another live-cell movie of actin in a COS-7 cell.

Supplementary Movie S3. A live-cell movie of ER in a U-2 OS Rtn4-Halo cell.

Supplementary Movie S4. A live-cell movie of mitochondria and lipid-droplet dynamics in a COS-7 cell with overlaid signal of OI-DIC and digital staining.

Supplementary Movie S5. An OI-DIC-only movie serving as the ground truth of Movie S4.

## Supporting information

A live-cell movie of actin in a COS-7 cell

Another live-cell movie of actin in a COS-7 cell

A live-cell movie of ER in a U-2 OS Rtn4-Halo cell

A live-cell movie of mitochondria and lipid-droplet dynamics in a COS-7 cell with overlaid signal of OI-DIC and digital staining

An OI-DIC-only movie serving as the ground truth of Movie S4

## Acknowledgements

We thank Dr. Maohan Su, Dr. Yuan Tian, Dr. Ed Courvan, Robby Nelson, Dr. Yang Li, and Dr. Hannahmariam Mekbib for insightful discussions on microscope design and cell biological applications. We thank Dr. Lukas Fuentes, Phylicia Kidd, and Dr. Yuan Tian for initial support setting up the experiments. We thank the Yale Center for Research Computing for guidance and use of the research computing infrastructure, specifically the McCleary cluster.

## Author contributions

Y.B, Z.M, M.S, and J.B. designed the study. Y.B. and Z.M. designed, built, and aligned the instrument. Y.B., Z.M. and D.B. extended the PYME software package for correlative OI-DIC and fluorescence imaging. X.C. and Q.L. wrote the machine learning algorithm. Y.B., X.C., and Q.L. trained the network. M.S. prepared the glass rod sample. Y.T. prepared the in vitro microtubule samples, S.S. and F.S. prepared the DNA origami samples, C.Z. prepared the Cajal Body samples, S.Y.S.F. and Y.B. prepared the spindle samples. Y.B. prepared all other samples, including the live-cell samples. Y.B. acquired all data and analyzed the data with input from other authors. K.M.N., J.H., C.L., D.B., M.S., and J.B. supervised the study. Y.B., Z.M., M.S., and J.B. wrote the manuscript with input from all authors.

## Data and materials availability

This study did not generate new reagents. All data needed to evaluate the conclusions in the paper are present in the paper, the Supplementary Materials, and the Movies. The PYthon Microscopy Environment (PYME) software package for the microscope hardware control and image post-processing is available in a public online repository (https://github.com/python-microscopy/python-microscopy). The deep learning algorithm is available in a public online repository (*will be added later*). The images are available on BioImage Archive (*will be added later*). Any additional data from this work can be obtained through the authors upon request.

## REFERENCES

1. D. Baddeley, J. Bewersdorf, Biological Insight from Super-Resolution Microscopy: What We Can Learn from Localization-Based Images. Annu. Rev. Biochem. 87, 965–989 (2018).

2. F. Zernike, How I discovered phase contrast. Science 121, 345–349 (1955).

3. W. Lang, Nomarski differential interference-contrast microscopy. (1982).

4. R. D. Allen, G. B. David, The Zeiss-Nomarski differential interference equipment for transmitted-light microscopy. Zeitschrift für Wissenschaftliche Mikroskopie und Mikroskopische Technik 69, 193–221 (1969).

5. R. Kasprowicz, R. Suman, P. O’Toole, Characterising live cell behaviour: Traditional label-free and quantitative phase imaging approaches. Int. J. Biochem. Cell Biol. 84, 89–95 (2017).

6. P. C. Chaumet, P. Bon, G. Maire, A. Sentenac, G. Baffou, Quantitative phase microscopies: accuracy comparison. Light Sci. Appl. 13 (2024).

7. G. Popescu, T. Ikeda, R. R. Dasari, M. S. Feld, Diffraction phase microscopy for quantifying cell structure and dynamics. Opt. Lett. 31, 775–777 (2006).

8. Y. Park, G. Popescu, K. Badizadegan, R. R. Dasari, M. S. Feld, Diffraction phase and fluorescence microscopy. Opt. Express 14, 8263 (2006).

9. Z. Wang, L. Millet, M. Mir, H. Ding, S. Unarunotai, J. Rogers, M. U. Gillette, G. Popescu, Spatial light interference microscopy (SLIM). Opt. Express 19, 1016–1026 (2011).

10. M. Mir, Z. Wang, Z. Shen, M. Bednarz, R. Bashir, I. Golding, S. G. Prasanth, G. Popescu, Optical measurement of cycle-dependent cell growth. Proc. Natl. Acad. Sci. U. S. A. 108, 13124–13129 (2011).

11. N. R. Subedi, P. S. Jung, E. L. Bredeweg, S. Nemati, S. E. Baker, D. N. Christodoulides, A. E. Vasdekis, Integrative quantitative-phase and airy light-sheet imaging. Sci. Rep. 10 (2020).

12. S.-M. Guo, L.-H. Yeh, J. Folkesson, I. E. Ivanov, A. P. Krishnan, M. G. Keefe, E. Hashemi, D. Shin, B. B. Chhun, N. H. Cho, Revealing architectural order with quantitative label-free imaging and deep learning. Elife 9, e55502 (2020).

13. I. E. Ivanov, E. Hirata-Miyasaki, T. Chandler, R. Cheloor-Kovilakam, Z. Liu, S. Pradeep, C. Liu, M. Bhave, S. Khadka, C. Arias, M. D. Leonetti, B. Huang, S. B. Mehta, Mantis: High-throughput 4D imaging and analysis of the molecular and physical architecture of cells. PNAS Nexus 3 (2024).

14. S. Shin, D. Kim, K. Kim, Y. Park, Super-resolution three-dimensional fluorescence and optical diffraction tomography of live cells using structured illumination generated by a digital micromirror device. Sci. Rep. 8 (2018).

15. S. Chowdhury, W. J. Eldridge, A. Wax, J. A. Izatt, Structured illumination microscopy for dual-modality 3D sub-diffraction resolution fluorescence and refractive-index reconstruction. Biomed. Opt. Express 8, 5776 (2017).

16. A. Descloux, K. S. Grußmayer, E. Bostan, T. Lukes, A. Bouwens, A. Sharipov, S. Geissbuehler, A.-L. Mahul-Mellier, H. A. Lashuel, M. Leutenegger, Combined multi-plane phase retrieval and super-resolution optical fluctuation imaging for 4D cell microscopy. Nat. Photonics 12, 165–172 (2018).

17. S. Chowdhury, W. J. Eldridge, A. Wax, J. A. Izatt, Structured illumination multimodal 3D-resolved quantitative phase and fluorescence sub-diffraction microscopy. Biomed. Opt. Express 8, 2496 (2017).

18. J. Ortega Arroyo, D. Cole, P. Kukura, Interferometric scattering microscopy and its combination with single-molecule fluorescence imaging. Nat. Protoc. 11, 617–633 (2016).

19. M. Shribak, R. Oldenbourg, Techniques for fast and sensitive measurements of two-dimensional birefringence distributions. Appl. Opt. 42, 3009–3017 (2003).

20. M. Shribak, Orientation-independent differential interference contrast microscopy technique and device, US Patent 7233434 (filed 17 December 2003).

21. M. Shribak, S. Inoué, Orientation-independent differential interference contrast microscopy. Appl. Opt. 45, 460–469 (2006).

22. M. Shribak, K. G. Larkin, D. Biggs, Mapping optical path length and image enhancement using quantitative orientation-independent differential interference contrast microscopy. J. Biomed. Opt. 22, 016006 (2017).

23. S. Iida, S. Ide, S. Tamura, M. Sasai, T. Tani, T. Goto, M. Shribak, K. Maeshima, Orientation-independent-DIC imaging reveals that a transient rise in depletion attraction contributes to mitotic chromosome condensation. Proc. Natl. Acad. Sci. U. S. A. 121 (2024).

24. Z. Marin, “Quantifying membrane topology at the nanoscale,” thesis, Yale University, New Haven, CT (2022).

25. J. E. Malamy, M. Shribak, An orientation-independent DIC microscope allows high resolution imaging of epithelial cell migration and wound healing in a cnidarian model. J. Microsc. 270, 290–301 (2018).

26. B. Huang, S. A. Jones, B. Brandenburg, X. Zhuang, Whole-cell 3D STORM reveals interactions between cellular structures with nanometer-scale resolution. Nat. Methods 5, 1047–1052 (2008).

27. A. E. S. Barentine, Y. Lin, E. M. Courvan, P. Kidd, M. Liu, L. Balduf, T. Phan, F. Rivera-Molina, M. R. Grace, Z. Marin, PYME: an integrated platform for high-throughput nanoscopy. Biophys. J. 121, 137a (2022).

28. R. Lin, A. H. Clowsley, T. Lutz, D. Baddeley, C. Soeller, 3D super-resolution microscopy performance and quantitative analysis assessment using DNA-PAINT and DNA origami test samples. Methods 174, 56–71 (2020).

29. J. Schnitzbauer, M. T. Strauss, T. Schlichthaerle, F. Schueder, R. Jungmann, Super-resolution microscopy with DNA-PAINT. Nat. Protoc. 12, 1198–1228 (2017).

30. S. Aknoun, J. Savatier, P. Bon, F. Galland, L. Abdeladim, B. Wattellier, S. Monneret, Living cell dry mass measurement using quantitative phase imaging with quadriwave lateral shearing interferometry: an accuracy and sensitivity discussion. J. Biomed. Opt. 20, 126009 (2015).

31. S. Chen, C. Li, Y. Zhu, Sensitivity evaluation of quantitative phase imaging: a study of wavelength shifting interferometry. Opt. Lett. 42, 1088–1091 (2017).

32. K. K. H. Chung, Z. Zhang, P. Kidd, Y. Zhang, N. D. Williams, B. Rollins, Y. Yang, C. Lin, D. Baddeley, J. Bewersdorf, Fluorogenic DNA-PAINT for faster, low-background super-resolution imaging. Nat. Methods 19, 554–559 (2022).

33. L. A. Fuentes, Z. Marin, J. Tyson, D. Baddeley, J. Bewersdorf, The nanoscale organization of reticulon 4 shapes local endoplasmic reticulum structure in situ. J. Cell Biol. 222 (2023).

34. D. Pirone, V. Bianco, L. Miccio, P. Memmolo, D. Psaltis, P. Ferraro, Beyond fluorescence: advances in computational label-free full specificity in 3D quantitative phase microscopy. Curr. Opin. Biotechnol. 85, 103054 (2024).

35. J. Park, B. Bai, D. Ryu, T. Liu, C. Lee, Y. Luo, M. J. Lee, L. Huang, J. Shin, Y. Zhang, D. Ryu, Y. Li, G. Kim, H.-S. Min, A. Ozcan, Y. Park, Artificial intelligence-enabled quantitative phase imaging methods for life sciences. Nat. Methods 20, 1645–1660 (2023).

36. N. Elmalam, L. Ben Nedava, A. Zaritsky, In silico labeling in cell biology: Potential and limitations. Curr. Opin. Cell Biol. 89, 102378 (2024).

37. C. Ounkomol, S. Seshamani, M. M. Maleckar, F. Collman, G. R. Johnson, Label-free prediction of three-dimensional fluorescence images from transmitted-light microscopy. Nat. Methods 15, 917–920 (2018).

38. Y. Jo, H. Cho, W. S. Park, G. Kim, D. Ryu, Y. S. Kim, M. Lee, S. Park, M. J. Lee, H. Joo, H. Jo, S. Lee, S. Lee, H.-S. Min, W. D. Heo, Y. Park, Label-free multiplexed microtomography of endogenous subcellular dynamics using generalizable deep learning. Nat. Cell Biol. 23, 1329–1337 (2021).

39. X. Chen, B. Zhou, L. Shi, H. Liu, Y. Pang, R. Wang, E. J. Miller, A. J. Sinusas, C. Liu, CT-free attenuation correction for dedicated cardiac SPECT using a 3D dual squeeze-and-excitation residual dense network. J. Nucl. Cardiol. 29, 2235–2250 (2022).

40. P. Marquet, C. Depeursinge, P. J. Magistretti, Review of quantitative phase-digital holographic microscopy: promising novel imaging technique to resolve neuronal network activity and identify cellular biomarkers of psychiatric disorders. Neurophotonics 1, 020901 (2014).

41. J. Kühn, F. Charrière, T. Colomb, E. Cuche, F. Montfort, Y. Emery, P. Marquet, C. Depeursinge, Axial sub-nanometer accuracy in digital holographic microscopy. Meas. Sci. Technol. 19, 074007 (2008).

42. P. Bon, G. Maucort, B. Wattellier, S. Monneret, Quadriwave lateral shearing interferometry for quantitative phase microscopy of living cells. Opt. Express 17, 13080–13094 (2009).

43. P. G. Kazansky, M. Shribak, Nanostructured birefringent optical elements and microscopes with nanostructured birefringent optical elements, European Patent Application EP (2023).

44. P. G. Kazansky, X. Chang, M. Shribak, “Using printed nanostructured optics with birefringence gradient in differential interference contrast (DIC) microscopy” (Focus on Microscopy, 2023).

45. Y. Sung, C. J. R. Sheppard, Three-dimensional imaging by partially coherent light under nonparaxial condition. J. Opt. Soc. Am. A Opt. Image Sci. Vis. 28, 554 (2011).

46. J. Roos, S. Bancelin, T. Delaire, A. Wilhelmi, F. Levet, M. Engelhardt, V. Viasnoff, R. Galland, U. V. Nägerl, J.-B. Sibarita, Arkitekt: streaming analysis and real-time workflows for microscopy. Nat. Methods 21, 1884–1894 (2024).

47. W. L. Stepp, E. B. Durmus, S. N. R. Alvarez, J. C. Landoni, G. Tortarolo, K. M. Douglass, M. Weigert, S. Manley, Smart hybrid microscopy for cell-friendly detection of rare events, bioRxiv (2025)p. 2025.04.04.647219.

48. O. M’Saad, M. Shribak, J. Bewersdorf, Unclearing Microscopy. bioRxivorg (2022).

49. C. Gell, V. Bormuth, G. J. Brouhard, D. N. Cohen, S. Diez, C. T. Friel, J. Helenius, B. Nitzsche, H. Petzold, J. Ribbe, E. Schäffer, J. H. Stear, A. Trushko, V. Varga, P. O. Widlund, M. Zanic, J. Howard, Microtubule dynamics reconstituted in vitro and imaged by single-molecule fluorescence microscopy. Methods Cell Biol. 95, 221–245 (2010).

50. Y. Tuna, A. Al-Hiyasat, J. Howard, Simultaneous interference reflection and total internal reflection fluorescence microscopy for imaging dynamic microtubules and associated proteins. J. Vis. Exp., e63730 (2022).

51. J. Schindelin, I. Arganda-Carreras, E. Frise, V. Kaynig, M. Longair, T. Pietzsch, S. Preibisch, C. Rueden, S. Saalfeld, B. Schmid, J.-Y. Tinevez, D. J. White, V. Hartenstein, K. Eliceiri, P. Tomancak, A. Cardona, Fiji: an open-source platform for biological-image analysis. Nat. Methods 9, 676–682 (2012).

52. D. Baddeley, D. Crossman, S. Rossberger, J. E. Cheyne, J. M. Montgomery, I. D. Jayasinghe, C. Cremer, M. B. Cannell, C. Soeller, 4D super-resolution microscopy with conventional fluorophores and single wavelength excitation in optically thick cells and tissues. PLoS One 6, e20645 (2011).

53. Y. Wang, J. Schnitzbauer, Z. Hu, X. Li, Y. Cheng, Z.-L. Huang, B. Huang, Localization events-based sample drift correction for localization microscopy with redundant cross-correlation algorithm. Opt. Express 22, 15982–15991 (2014).

