## Supplementary Materials for "A correlative quantitative phase contrast and fluorescence super-resolution microscope for imaging molecules in their cellular context"

**This PDF file includes:**

Figs. S1 to S4

**Other Supplementary Materials for this manuscript include the following:**

Movies S1 to S5

**Supplementary Figures**

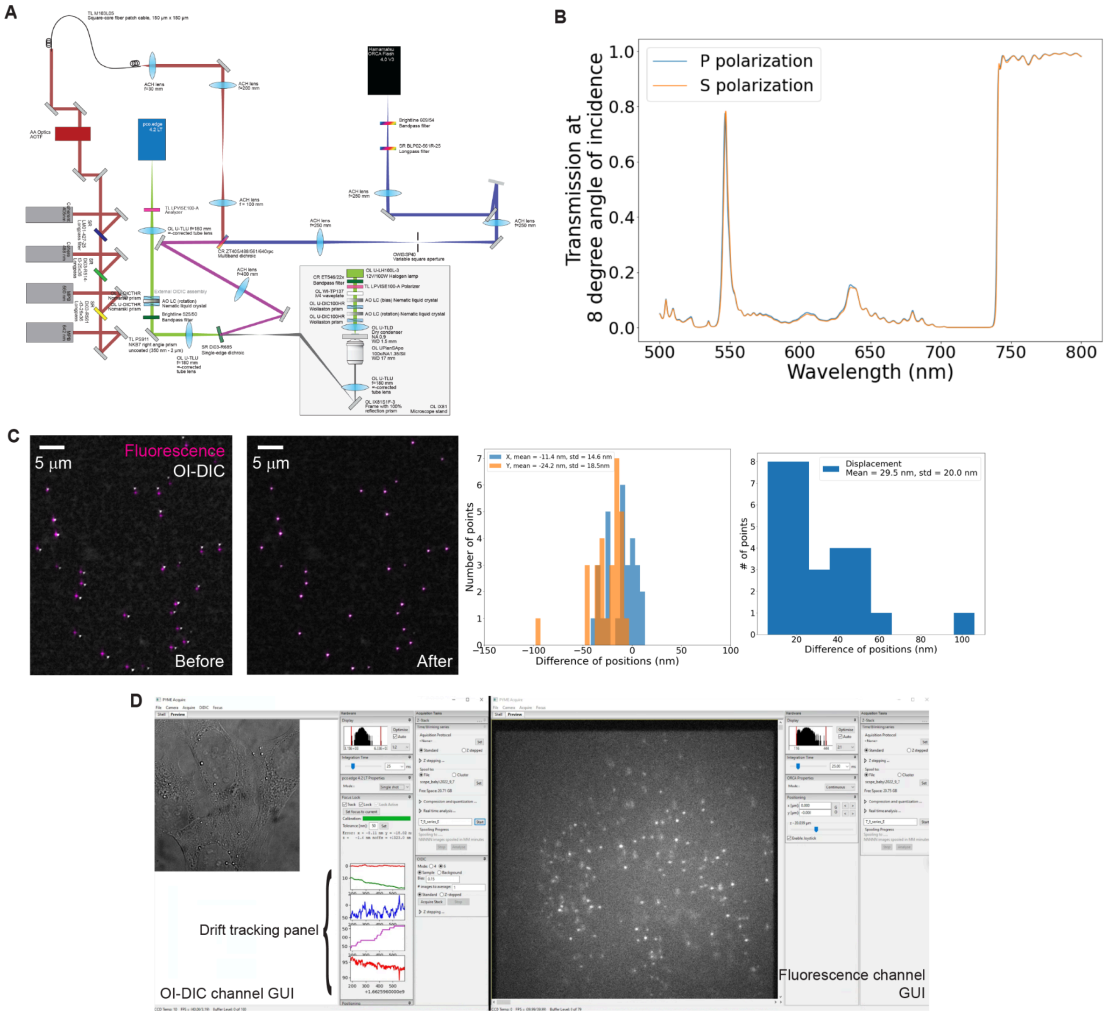


**Fig. S1. Optical setup and characterization of the correlative microscope.** (**A**) A detailed schematic of the correlative OI-DIC/SMLM microscope. (**B**) Transmission spectrum of the dichroic (Semrock, Di03-R685) at 8° angle of incidence, showing the spectral difference between s and p polarization is ignorable. (**C**) Shiftmap calibration using 200-nm-diameter beads. Before the calibration, the overlaid fluorescence and OI-DIC image of 200-nm-diameter beads shows obvious displacement (left); after the calibration, the overlaid image shows overlapping fluorescence and OI-DIC signal (middle); the remaining displacement after the shiftmap calibration is shown in the histogram and is below the diffraction limit (right). (**D**) Graphical user interface of simultaneous software-based drift tracking (left) and SMLM acquisition (right). During the drift tracking, the system automatically corrects the focal plane displacement by moving the piezo and records lateral drift. The four plots in the drift tracking panel, from top to bottom, show lateral drift in x and y, piezo position after the focal plane correction, total focal plane displacement before correction, and correlation factor between the live-view and calibrated images, respectively.

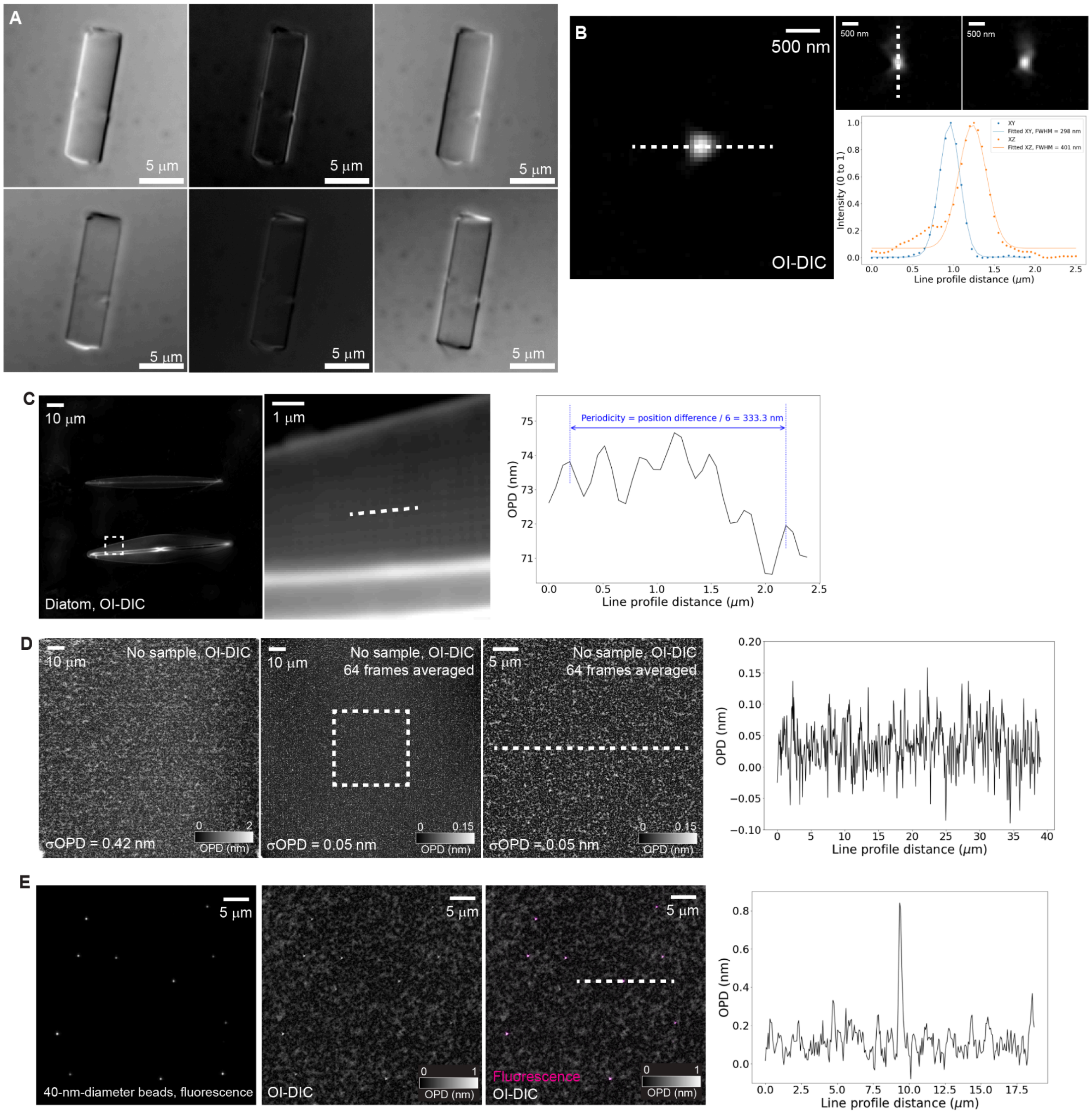


**Fig. S2. Characterization of the OI-DIC channel.** (**A**) Six raw DIC images of the 4.1-µm-diameter glass rod for characterizing quantitative measurement of the OI-DIC channel. (**B**) OI-DIC PSF calibration by imaging 100-nm-diameter beads in the OI-DIC channel, showing 298 nm lateral resolution and 401 nm axial resolution. (**C**) OI-DIC lateral resolution validation by imaging diatom *F. rhomboides*. An OI-DIC image of diatom *F. rhomboides* (left). The striae are visible in the magnified view of the dashed box (middle), and the line profile along the dashed line shows the distance between striae is about 333.3 nm, exceeding the OI-DIC lateral resolution limit by 12%. (**D**) OI-DIC sensitivity assessment. The sensitivity is defined by the smallest OPD that can be detected by the OI-DIC, which is quantified by the standard deviation of OPD measurements without any sample on the microscope. Without averaging acquisitions, the OI-DIC has a sensitivity of ~0.42 nm; by averaging 64 acquisitions, it achieves a sensitivity ~0.05 nm. (**E**) Correlative fluorescence and OI-DIC imaging of 40-nm-diameter beads, demonstrating the high sensitivity of the OI-DIC channel.

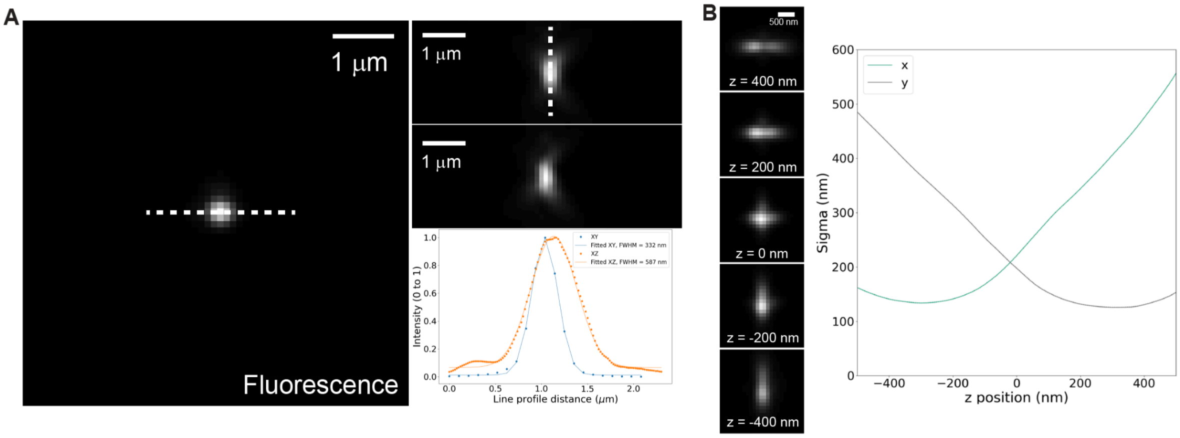


**Fig. S3. PSF calibration of the fluorescence channel.** (**A**) Diffraction-limited fluorescence PSF calibrated by imaging 100-nm-diameter beads, showing 332 nm lateral resolution 587 nm axial resolution. (**B**) Astigmatic fluorescence PSF calibrated by imaging 100-nm-diameter beads, with a cylindrical lens implemented near the intermediate image plane of the fluorescence channel.

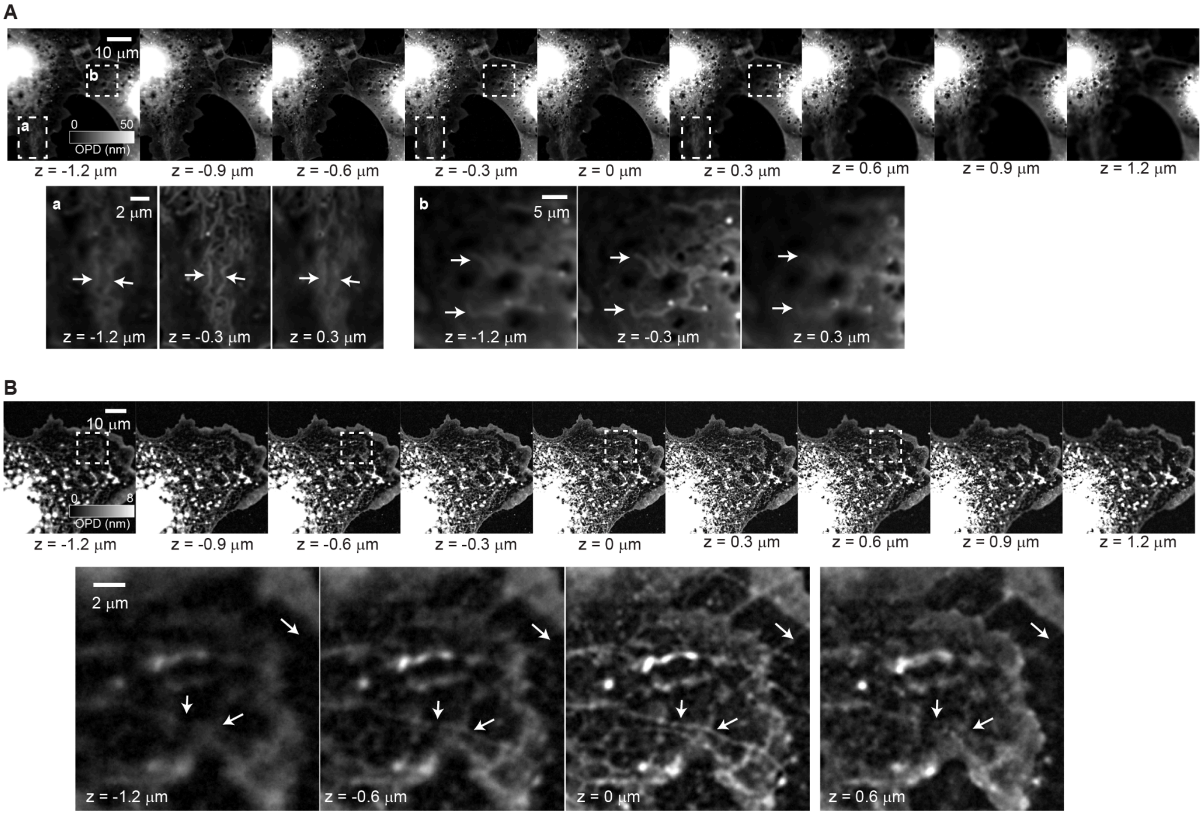

**Fig. S4. Example of visualizing mitochondria and microtubules in unlabeled fixed COS-7 cells in OI-DIC with z-stacking.** (**A**) Visualization of mitochondria in unlabeled fixed COS-7 cells with z-stacking. Arrows in the magnified regions of the dashed boxes highlight mitochondria that transit from out of focus to in focus and back to out of focus during z-stacking. (B) Visualization of microtubules in unlabeled fixed COS-7 cells with z-stacking. Arrows in the magnified regions of the dashed boxes highlight microtubules that transit from out of focus to in focus and back to out of focus during z-stacking.
